# A scalable, Rotating Disc Bioelectrochemical Reactor (RDBER) suitable for the cultivation of both cathodic and anodic biofilms

**DOI:** 10.1101/2022.09.12.507646

**Authors:** Max Hackbarth, Johannes Gescher, Harald Horn, Johannes Eberhard Reiner

**Affiliations:** Engler-Bunte-Institut, Water Chemistry and Water Technology, Karlsruhe Institute of Technology (KIT), Engler-Bunte-Ring 9a, 76131 Karlsruhe, Germany; Karlsruhe Institute of Technology (KIT), Institute for Applied Biosciences (IAB), Fritz-Haber-Weg 2, 76131, Karlsruhe, Germany; Karlsruhe Institute of Technology (KIT), Institute for Biological Interfaces (IBG 1), Hermann-von-Helmholtz-Platz 1, 76344 Eggenstein-Leopoldshafen, Germany; Hamburg University of Technology, Institute of Technical Microbiology, Kasernenstraße 12, 21073 Hamburg, Germany; DVGW Research Centre, Water Chemistry and Water Technology, Engler-Bunte-Ring 9a, 76131 Karlsruhe, Germany

**Keywords:** Pure culture microbial electrosynthesis, microbial electrolysis cell, *Kyrpidia spormannii*, rotating disc bioelectrochemical reactor, rotating biological contactor

## Abstract

This study discusses the construction and operation of a membrane-less bioelectrochemical reactor that employs rotating working electrodes with a surface area of up to 1 m^2^. As a proof-of-principle for an aerobic microbial electrosynthesis process, *Kyrpidia spormannii* was cultivated in the reactor. Optical coherence tomography was used to examine the spatial distribution of the cathodic biofilm. After 24 days 87% of the cathode surface was covered with biofilm that was characterized by a radial increase in its biovolume towards the circumcenter of the electrodes reaching up to 92.13 μm^3^ μm^-2^. To demonstrate the versatility of the system, we further operated the reactor as a microbial electrolysis cell employing a co-culture of *Shewanella oneidensis* and *Geobacter sulfurreducens*. Anodic current densities of up to 130 μA cm^-2^ were measured during these batch experiments. This resulted in a maximum production rate of 0.43 liters of pure hydrogen per liter reactor volume and day.

**Graphical Abstract:** 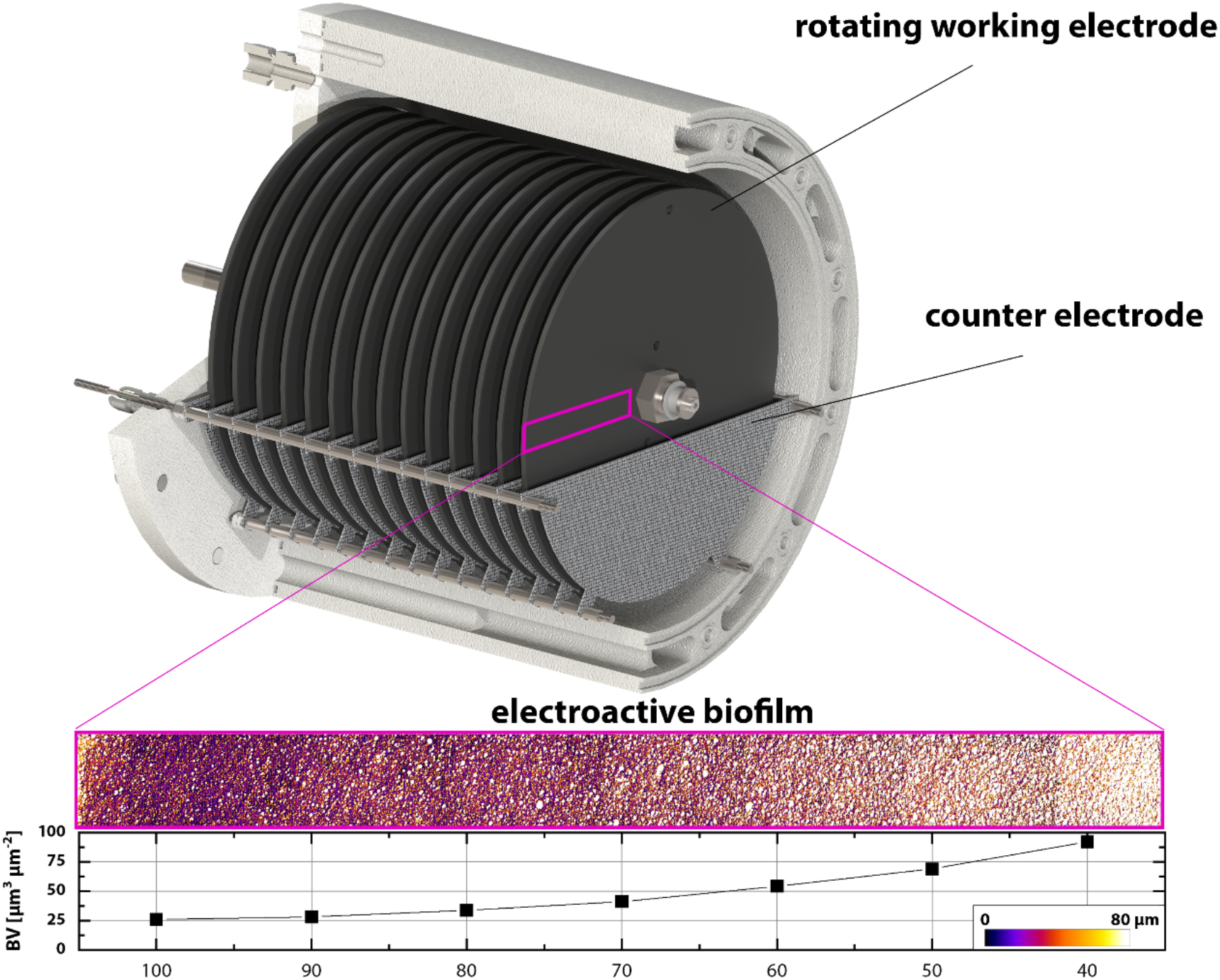

**Highlights:**

- Construction of a 10 L membrane-less, pressurizable bioelectrochemical reactor
- Rotating working electrodes with up to 1 m^2^ electrode surface
- Electroautotrophic cultivation and quantification of *K. spormannii* biofilms
- Initial cell density crucial for successful *K. spormannii* biofilm formation
- Anodic operation as MEC with *Shewanella* / *Geobacter* coculture

## 1. Introduction

Tapering off fossil resources by renewable commodities that are based on a circular utilization of resources is key to the transition towards a sustainable economy (Lieder and Rashid, 2016). In this context, future (bio)chemical transformations of raw materials and waste streams must be fueled by renewable electrical energy (Krieg et al., 2018a). The so-called microbial electrosynthesis (MES) is a promising biotechnological example for such a conversion of electricity and carbon dioxide into value-added organic compounds (Nevin et al., 2010). This process relies on electroactive microorganisms that serve as a biocatalyst on the cathode of a bioelectrochemical system (BES) (Logan et al., 2019). By applying a voltage to the BES, typically water is oxidized at the anode into oxygen and protons. Driven by a power source the released electrons are subsequently used either directly or indirectly by the autotrophic, cathodic biofilm as the sole energy and electron source for the reduction of CO_2_. Although MES could theoretically outperform abiotic carbon utilization technologies as well as established gas fermentations in some specific applications, the technology is not yet economically feasible on a large scale (Christodoulou and Velasquez-Orta, 2016; Jourdin et al., 2020; Jourdin and Burdyny, 2021; Prévoteau et al., 2020; Wood et al., 2021). However, the novel electroautotrophic species *Kyrpidia spormannii* EA-1^T^ was isolated from a thermophilic cathodic community recently (Reiner et al., 2018a, 2018b). This thermoacidophilic ‘Knallgas’ bacterium is a natural producer of polyhydroxyalkanoates (PHAs) and was shown to thrive on cathodes as the sole electron and energy source for the reduction of CO_2_ (Reiner et al., 2020). Hence, *K. spormannii* is a promising candidate for MES - not only since PHAs are precursors for biodegradable bioplastics - but also due to the organisms comparably good biofilm-formation abilities on cathodes. Thus, it was shown that *K. spormannii* forms robust biofilms with a thickness of well over 100 μm – an important prerequisite for MES as stated recently by Vassilev and colleagues (Barham et al., 1984; Hackbarth et al., 2020; Vassilev et al., 2022). Although it seems to be an intrinsic property of electroautotrophic organisms, the molecular details of extracellular electron uptake (EET) from a cathode are also subject to keen debate with regard to *K. spormannii* (Jung et al., 2020; Pillot et al., 2022a; Prévoteau et al., 2020; Reiner et al., 2020).

While - with respect to the inanimate components of a MES system - much effort has been put into studying the effect of material and structural properties of electrodes, the development of new and, in particular, scalable reactor concepts for MES has been rather sidelined (Enzmann et al., 2019b; Guo et al., 2015; Kerzenmacher, 2017; Pillot et al., 2022a, 2022b; Zhang et al., 2021). This is in spite of the fact that whether microbial electrochemical technologies (METs) can carve a niche in a future sustainable economy depends strongly on the availability of a suitable reactor technology. Typical reactor systems for the cultivation of electroactive microorganisms in laboratory settings are cube, bottle, or flat-plate-type reactors (Enzmann et al., 2019a; Krieg et al., 2018b). Especially for MES applications, a semipermeable membrane is commonly required to spatially separate anode and cathode compartments. Membranes can be advantageous when extremely high Coulombic efficiencies are desired, as they avoid chemical short circuits caused by diffusion of electrochemically active chemicals (Krieg et al., 2018b). Moreover, due to anodically produced oxygen a membrane is particularly crucial if strict anaerobes, such as acetogenic or methanogenic organisms, serve as cathodic biocatalysts for MES (Gildemyn et al., 2017). However, the utilization of membranes leads to significantly increased surface-specific ohmic losses as a plain result of an increased distance between anode and cathode, but also through diffusion limitations often caused by hardly mixed zones around the membranes. The latter may also result in undesirable pH shifts in the separated compartments (Gildemyn et al., 2017; Guo et al., 2017; Krieg et al., 2018b; Prévoteau et al., 2020; Zeppilli et al., 2016). Finally, most membranes are not autoclavable (according to the manufacturer), sensitive to fouling and prone to leakage due to lack of mechanical stability (Utesch and Zeng, 2018). The above-mentioned disadvantages of membranes, along with the resulting complexity of the reactor set-up, lead to a high-maintenance reactor system which is likely to be very cost-intensive in long-term operation (Kracke et al., 2018).

Here, we present the concept of a membrane-less, scalable rotating disc bioelectrochemical reactor (RDBER) for aerobic MES and other pure culture bioelectrochemical applications. The reactor is fully autoclavable and can be pressurized to at least 0.5 bar. Moreover, it is equipped with an optical window allowing for in vivo monitoring of biofilm development on the electrodes using optical coherence tomography (OCT). The systems design is inspired by a so-called rotating biological contactor (RBC) (Hassard et al., 2015). RBC is a fixed-film bioreactor used in wastewater treatment that consists of closely spaced discs that are mounted on a common horizontal shaft and are typically partially submerged in liquid. Cortez and colleagues highlight the low maintenance and operating costs, ease of process control and monitoring, and low land requirements as particular advantages of RBC (Cortez et al., 2008). This is mainly due to the fact that the flow velocity across the biofilm enhancing substrate supply is not achieved by energy-intensive pumping of high volumes or stirring, but by rotation of the biofilm-covered discs. The first steps in the “electrification” of RBCs have already been successfully carried out on a laboratory scale (Cheng et al., 2012, 2011; Rodziewicz et al., 2019; Sayess et al., 2013): Several attempts have been made to use conductive rotating discs as cathode or anode of a bioelectrochemical system. However, these studies consistently focus on the purification of wastewater. Hence, biotechnological requirements for a pure culture MES process (e.g., working with pure cultures under sterile conditions, control of temperature and pH, or pressurizability to increase the solubility of the gaseous substrates like CO_2_) were not considered in the reactor design.

The RDBER operates in either anodic (as a microbial electrolysis cell) or cathodic mode with an effective working electrode area of up to 1 m^2^ (depending on the number of graphite discs mounted). As a proof of principle for an axenic, aerobic MES-process we cultivated *K. spormannii* in the system and correlated the OCT-derived, spacial resolved biovolume of the cathodic biofilm with the radius of the rotating cathodes. Furthermore, we demonstrated that the developed system can also be operated as a microbial electrolysis cell (MEC). In a MEC, electroactive organisms oxidize organic matter into CO_2_ and protons, whereby the released electrons are transferred from the organisms to the anode, which serves as a terminal respiratory electron acceptor. From the anode, the electrons are then transferred to the counter electrode (cathode) by means of a voltage source, in order to reduce protons abiotically into hydrogen. For MEC-operation of the RDBER a co-culture of *Shewanella oneidensis* and *Geobacter sulfurreducens* was cultivated as an anodic biofilm on the rotating working electrodes. Both organisms are well-studied electroactive model organisms for extracellular electron transfer of respiratory electrons to insoluble electron acceptors such as an anode of a MEC (Edel et al., 2022; Korth and Harnisch, 2019; Logan et al., 2019).

Thus, by cultivating defined pure cultures in different operating modes (anodic/cathodic) in each case, we were able not only to demonstrate the general functionality of the system, but also to provide a first insight into the range of possible applications offered by the RDBER as a technological platform for METs.

## 2. Material and methods

### 2.1 Reactor Design

The design of the reactor system is illustrated in detail in Fig. 1. The reaction chamber, consisting of the reactor jacket (component b in Figs. 1A and 1C) and two flanges, was designed as a horizontal cylinder made of polypropylene. The inner diameter of the jacket measuring 238 mm and the length of 230 mm result in an inner volume of the reaction chamber of 10 L. An inspection window made of polycarbonate (Fig. 1A, component a) was integrated in the front flange allowing the observation of the foremost working electrode (Figs. 1A and 1C, component d). A circular recess milled in the center of the window serves as a guide and seat for the shaft bearing (component f in Fig. 1A). Figure 1C shows a cross-sectional view of the RDBER illustrating the arrangement of the electrodes in the reaction chamber. The shaft, which was made of high-purity titanium and sealed from the environment by a shaft seal, is located concentrically in the reactor. A precisely aligned shaft bearing in the rear flange, together with the fixed bearing in the front window, ensures that the shaft runs as friction-free and balanced as possible. For compensation of the temperature-related expansion of the shaft relative to the reaction chamber, the bearing in the rear flange was designed as a floating bearing. The shaft was manufactured as a threaded rod onto which working electrode holders (custom made titanium nuts) are mounted. The working electrode discs made of graphite are clamped between two holders in each case and centered in the process. This ensures that the discs are in contact with the shaft and thus also with the voltage source. In total, up to 14 working electrode discs with a thickness of 3 mm and a diameter of 210 mm can be mounted on the shaft. This corresponds to an active working electrode surface (front and back) of up to 1 m^2^ and, relative to the reaction chamber volume, a surface-to-volume ratio of 100 m^2^ m^-3^.

**Fig. 1:**
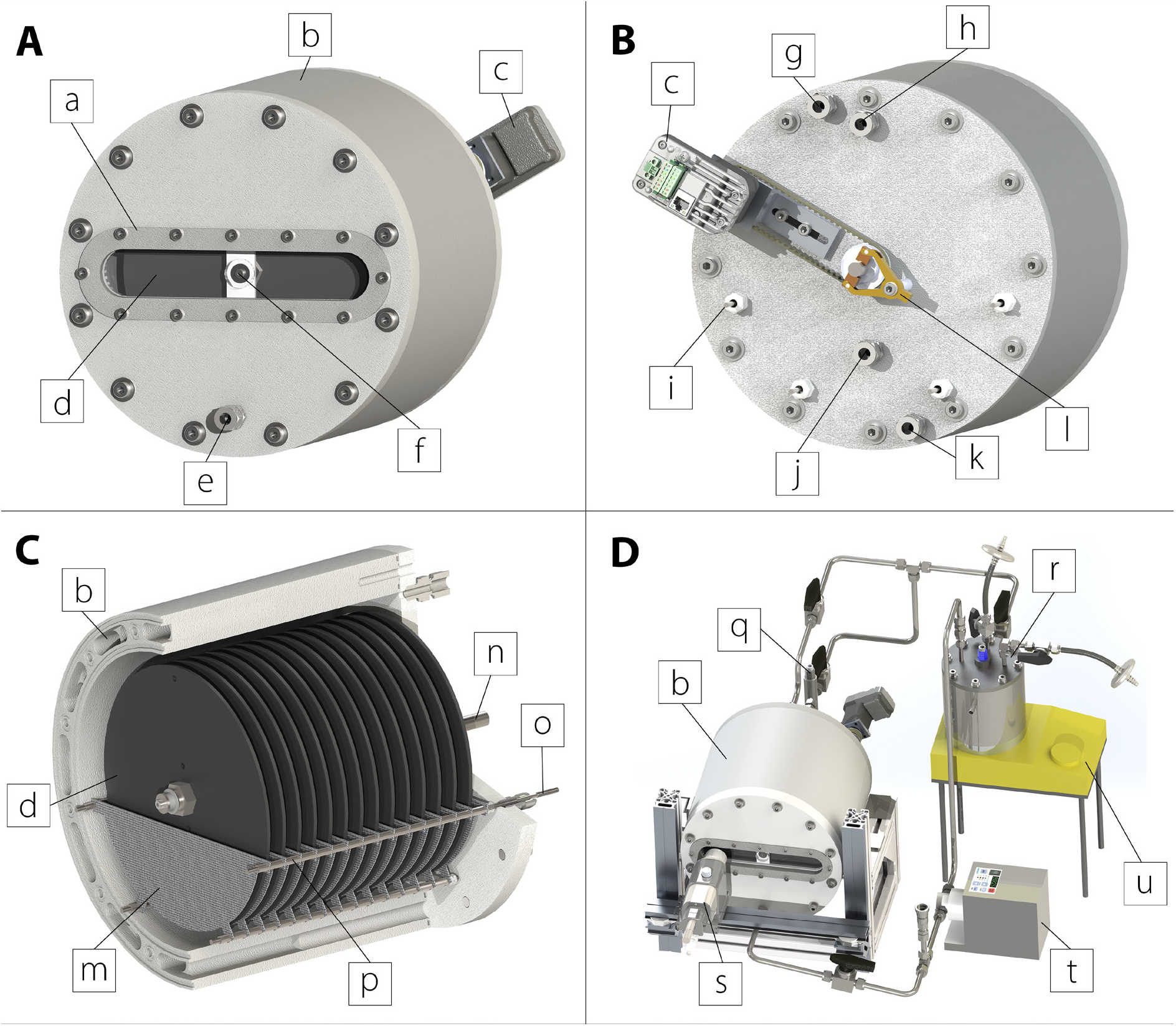
3D renderings of the RDBER: (A) Front view of the reaction vessel, (B) rear view of the reaction vessel, (C) cross-section of the reaction vessel, (D) full system with stainless-steel vessel and piping. (a – inspection window, b – reactor jacket, c – stepper motor, d – working electrode disc, e – inlet to reaction chamber, f – shaft journal bearing, g – outlet of heating jacket, h – gas outlet, i – connector to counter electrode, j – outlet of reaction chamber and connection to reference electrode holder, k – inlet to heating jacket, l – carbon brush holder as connector to rotating working electrodes, m – counter electrodes, n – working electrode shaft, o – counter electrode shaft, p – counter electrode spacers, q – reference electrode, r - double jacketed vessel with integrated pH electrode, s – OCT probe head, t – magnetic gear pump, u – magnetic stirrer).

Since in RBC-like systems the cross-flow across the biofilm is adjusted by the rotational speed of the rotating substrate (here the working electrode), a high-resolution stepper motor (component c, Figs. 1A and 1B) was implemented (Lexium MDrive Motion Control, Schneider Electric) on the rear flange. The employment of such a stepper motor allows, in addition to the load-independent control of the rotational speed, the position-accurate visualization of selected locations on the substrate. Apart from the toothed belt wheel required for this purpose, a double leg carbon brush holder (Fig. 1B, component l) with silver-graphite brushes was attached to the end of the titanium shaft (Fig. 1C, component n). Via this brush holder, electrical contact was made from the potentiostat to the rotating shaft and thus also to the working electrodes. The counter electrodes were made of titanium sheet with an iridium mixed oxide coating (platinode type 177, Umicore) in the form of half-rounds (Fig. 1C, component m) and connected to each other by means of four titanium threaded rods (Fig. 1C, component o). Exactly cut pieces of titanium pipe (component p in Fig. 1C) are mounted between each two anode sheets to provide the desired distance between the working and counter electrodes. The distance between anode and cathode amount to 5.5 mm. A total of up to 15 fixed counter electrode sheets with an active area of approx. 0.5 m^2^ can be installed in the reaction chamber in this way. The four titanium threaded rods are led out of the reaction chamber through the rear flange (Fig. 1B, component i) and can thus be connected to the potentiostat. Cable bushings made of polytetrafluoroethylene (PTFE) provide the necessary sealing.

In order to supply the growing biofilm with CO_2_-enriched media during cathodic operation, the gas-saturated and pre-tempered culture medium was pumped by means of a gear pump with an autoclavable pump head (REGLO-Z Digital, Z-1830 P, Ismatec) (Fig. 1D, component t) from an optional stainless-steel (component r in Fig. 1D) double jacket container via a compression fitting (Fig. 1D, component e) located in the front flange into the reaction chamber. The medium was then circulated in the overall system via two screw fittings (parts h and j in Fig. 1B) located on the rear flange. During operation the volume flow was led via the compression fitting j (Fig. 1B) connected to the reference electrode holder containing an Ag/AgCl reference electrode (part q in Fig. 1D) (SE23I, Meinsberg Sensortechnik). The second screw connection h (Fig. 1B) served as an outlet for any gas that may have formed and collected in the reaction chamber.

To ensure constant temperatures in the reaction chamber, heating channels were milled into the 31 mm thick polypropylene jacket. These channels were flushed with heating liquid and are shown in the cross-section of the reaction chamber (Fig. 1C). Inlet k and outlet g (Fig. 1B) of the heating jacket were placed at the rear flange of the reaction chamber and coupled to a circulation thermostat via clamping ring connections. Simultaneously the growth medium was heated supportively in the optional stainless-steel double jacket vessel during cathodic operation (Fig. 1D, part r). Moreover, the gaseous substrates for electroautotrophic growth were continuously passed through the headspace of the stainless-steel vessel under pressure. The inlet and outlet of the substrate gases are equipped with sterile filters to prevent contamination. By installing a pH electrode (Ceragel CPS71D, Endress + Hauser) in the vessel, the pH value and temperature was logged and the circulation thermostat was regulated. The RDBER was aligned and rigidly fixed on a base frame. Moreover, a holder on which the OCT probe (component s in Fig. 1D) can be fixed was mounted to the base frame.

### 2.2 Strains and preculture conditions

Electroautotrophic experiments were conducted with a population of *K. spormannii* EA-1^T^ previously adapted to cathodic growth (Jung et al., 2020). The preculture for RDBER experiments was grown from freezer stocks in a modified R2A medium (yeast extract 5 g L^-1^, tryptone 1 g L^-1^, casamino acids 0.5 g L^-1^, sodium pyruvate 0.5 g L^-1^, MgSO_4_·7 H_2_O 0.05 g L^-1^, K_2_HPO_4_ 0.1 g L^-1^, MOPS 10 mM - pH 6) at 60 °C without agitation. The heterotrophically grown pre-culture of *K. spormannii* was washed two times with ES-medium (pH 3.5; 10 mM NH4Cl, 2.5 mM NaCl, 0.3 mM KH_2_PO_4_, 0.12 mM MgSO_4_, 0.1 mM CaCl_2_, 1 mL L^-1^ Wolfe’s mineral elixir) before inoculating the autoclaved RDBER setup (Reiner et al., 2018a). Depending on the desired planktonic cell number per working electrode surface ratio the initial optical density (OD_600_) in the RDBER was 0.1 or 0.3 respectively. A counting chamber (Bürker-Türk) was consulted in order to determine the specific cell number per OD_600_ and Volume (an OD_600_ of 1 corresponded to 3.16±0.25 * 10^7^ *K. spormannii* cells per mL).

*A* mixed culture of the model organisms *Shewanella oneidensis* MR-1 and *Geobacter sulfurreducens* PCA was employed as biocatalyst during anodic operation of the RDBER. *S. oneidensis* was precultured aerobically in LB-Medium (Lennox), while the cultivation of the *G. sulfurreducens* preculture was conducted under anoxic conditions in BES-Medium [0.42 g L^-1^ KH_2_PO_4_, 0.22 g L^-1^ K_2_HPO_4_, 0.2 g L^-1^ NH4Cl, 0.38 g L^-1^ KCl, 0.36 g L^-1^ NaCl, 0.21 g L^-1^ MgCl_2_, 11.92 g L^-1^ HEPES – after autoclaving the medium was complemented with 1 mL L^-1^ 0.4 mM CaCl_2_ solution, 10 mL L^-1^ NB trace mineral solution, 1.0 mL selenite–tungstate solution (13 mM NaOH, 17 μM Na_2_SeO_3_, and 12 μM Na_2_WO_4_), 10 mL vitamin solution (German Type Culture Collection, DSMZ, media 141) and then sparged in rubber sealed bottles with N_2_/CO_2_ (80:20 vol%) for 30 minutes]. For preculturing, the BES-medium was complemented with 20 mM sodium acetate as electron donor and disodium fumarate (40 mM) as electron acceptor. The final pH was set to 7.4. Before inoculating the reactor with a mixture of *G. sulfurreducens* and *S. oneidensis* in a 9 to 1 ratio to an initial OD_600_ of 0.1, the cells were washed two times in BES-Medium.

### 2.3 Cultivation of cathodic *K. spormannii* biofilms

After pumping the previously inoculated ES-Medium omitting any organic carbon source (Reiner et al., 2020) into the sterile RDBER, the medium temperature was adjusted to 60 °C. The headspace of the mixing vessel was pressurized to 0.5 bar and continuously flushed with a mixture of 99.5% CO_2_ (v/v) and 0.5% O_2_ (v/v) while the liquid phase was continuously stirred at 300 rpm. The additional stirring of the medium in the mixing vessel by means of a magnetic stirrer (see component u in Fig. 1D) ensured improved mass transfer of the gases and increased the heat transfer of the heated jacket into the medium. The cultivation medium enriched with the gaseous substrates was pumped through the RDBER at a constant volumetric flow rate of 500 mL min^-1^. In order to achieve different planktonic cell number per working electrode surface ratios, either only one or all electrodes were mounted (714 cm^2^ vs. 1 m^2^ working electrode surface). The rotational speed of the working electrodes amounted to one revolution per minute. A constant current density of 1 μA cm^-2^ was applied for 24 h immediately after inoculation by means of a potentiostat (Interface 5000P, Gamry Instruments, Warmister, USA). After this initial polarization phase, the current density was set to 3 μA cm^-2^ by the end of the experiment. The experiments were stopped upon detection of no further biofilm increase.

### 2.4 Anodic operation as Microbial Electrolysis Cell

10 L of inoculated BES-medium containing 20 mM sodium acetate and 20 mM sodium lactate as carbon source was pumped in the previously autoclaved RDBER. Chronoamperometric batch experiments were performed at 30 °C and a constant working electrode potential of 0 mV vs. SHE (standard hydrogen electrode) was set. For anodic operation the stainless-steel mixing vessel was omitted while the pipeline was shorted with a pH-electrode holder. Evolving reaction gases (e.g. Hydrogen and CO_2_) were collected at the upper outlet h (see Fig. 1B) and led through a MilliGascounter (Ritter, Germany) into a gas sampling bag. Analysis of collected gases was conducted by means of a Micro GC (490 Micro GC, Agilent Technologies, Germany) using two different carrier gases (argon and helium) and two stationary phases (CP-Molsieve 5 Å Plot and PoraPLOT U J&W GC columns, Agilent Technologies, Germany). Moreover, liquid samples were taken twice a day in order to quantify the concentration of organic acids by means of a Metrohm 881 Compact Pro Ion Exchange Chromatograph with a Metrosep Organic Acids 250/7.8 column (Metrohm, Switzerland).

### 2.5 OCT imaging and digital image processing

A GANYMEDE II spectral domain OCT system (Thorlabs GmbH, Dachau, Germany) equipped with a LSM04 objective lens was employed for 3D biofilm visualization (Thorlabs GmbH, Dachau, Germany). Prior to each OCT acquisition, the stepper motor was consistently halted in the same pre-assigned position. Together with the fixed position of the OCT probe holder, this allowed reproducible visualization of a selected radial cathode section. OCT 3D data sets (C-scans) were acquired regularly over the visualizable radial area (see Fig. 2). Each spot was scanned with a size of 10.0 mm × 6.0 mm × 2.0 mm (L × W × H) and pixel resolutions of 8 μm px^-1^ (lateral, xy-plane) and 2.06 μm px ^-1^ (axial, z-direction). Adjacent OCT data sets were then stitched to depict the whole radius of the cathode. The generated OCT datasets were then analyzed, merged, and finally assembled into a height profile (6 mm × 70 mm; H × W). The visualized rectangular area was then trimmed into an isosceles trapezoid for final evaluation. The trapezoid represents a circular sector of the round disc, which abuts the outer radius of the substrate and was shortened to 70 mm by cutting off the tip. 110 of these trapezoids could thus be approximately combined to form a circular ring with an outer diameter of 21 mm and an inner diameter of 70 mm. Finally, the mean biovolume 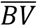, the mean accumulation rate 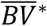, and the mean coverage 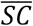 over the radius were determined from the cropped trapezoid (detailed view Figure 2). In addition, the trapezoid was divided into 7 further 10 mm long sub-trapezoids to obtain detailed information about the spatial distribution of biovolume (*BV*) along the radius of the circular blank. ImageJ (Schneider et al., 2012), operated with in-house made macros (Bauer et al., 2019), was used to calculate topographic representations of the bulk-biofilm interface (height maps) and the structural parameters used for biofilm characterization.

**Fig. 2:**
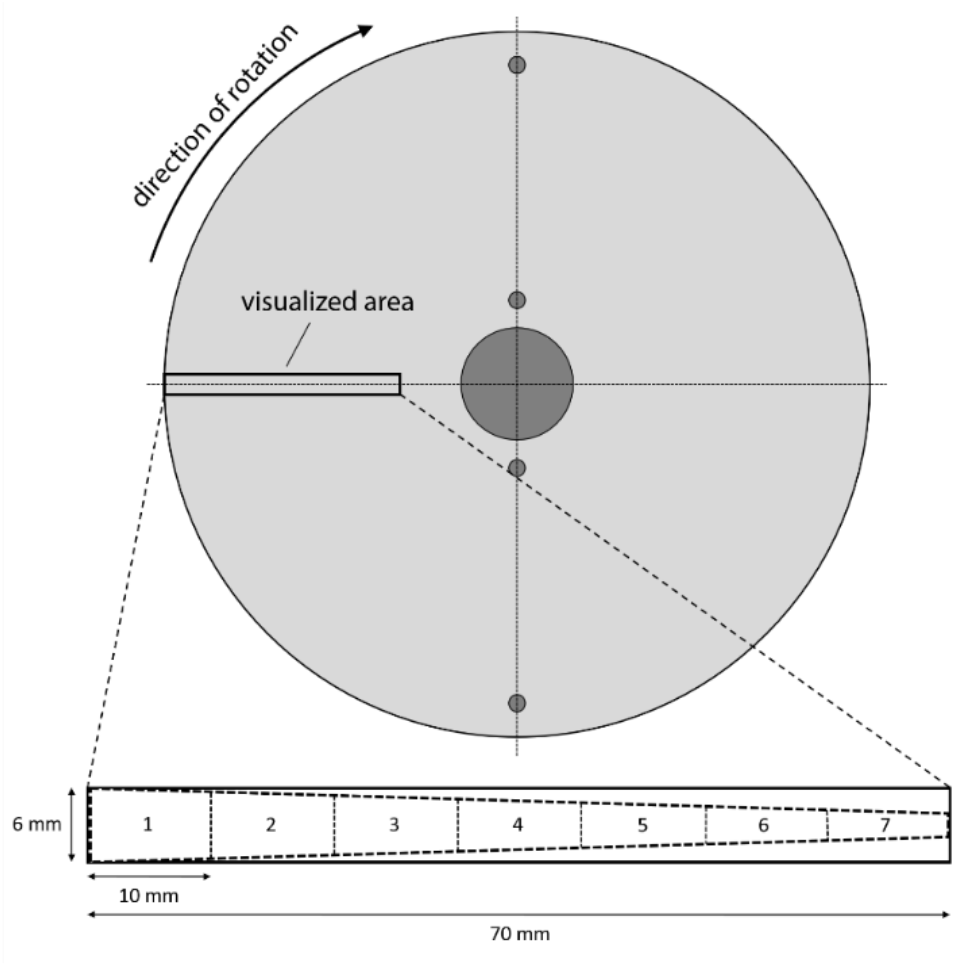
Representation of the visualized area and the analyzed trapezoidal sections of the generated OCT data sets.

### 2.6 Parameters for biofilm characterization

The calculation of the biofilm structure parameters - biovolume *BV* (μm^3^ μm^-2^), mean biovolume 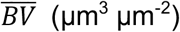, mean biofilm accumulation rate 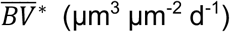 and mean substratum coverage 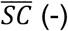, was based on height maps according to Wagner and Horn(Wagner and Horn, 2017).

Biovolume *BV* is defined as the measured biofilm volume *V_BF_* (μm^3^) per specific area of a single small trapezoid *A_ST_* (μm^2^)

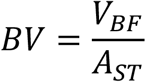

whereas 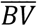 is being defined as

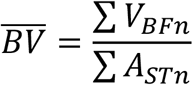

Where *n* indexes the individual small trapezoids.

Mean biofilm accumulation rate 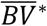 corresponds to 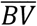 derived by time:

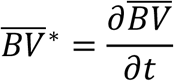

Finally, mean substratum coverage 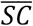 is described as

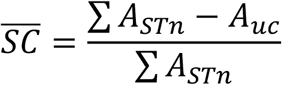

with the area of uncovered substratum *A_uc_* (biofilm thickness = 0 μm).

## 3. Results and discussion

### 3.1 Cultivation of cathodic *K. spormannii* biofilms

In order to prove the functionality of the newly developed reactor concept, two different pure culture cultivations of electroactive microorganisms were performed in the RDBER. Moreover, initial cultivation experiments performed with *K. spormannii* were designed to answer the question of whether an RBC-like reactor could even be a potential technological platform for upscaling a biofilm-based, aerobic MES process that was previously demonstrated in small-scale, custom-made flow cells (Hackbarth et al., 2020; Jung et al., 2020). Thus, as many process parameters as possible were transferred from the established flow cell cultivation to the cultivation of *K. spormannii* in the RDBER for a preliminary cultivation experiment (see Table 1). After several days of reactor operation, OCT analysis of the cathode revealed isolated tower-like structures formed on the cathode. However, their quantity and height decreased sharply again in the course of a few days (Fig. S1). Moreover, the surface coverage of the substratum never exceeded 20% even after one month of cultivation. In summary, no continuous biofilm could be detected on the cathodes by OCT during the preliminary cultivations. When examined more closely, there were two particular process parameters that deviated from the flow cell experiments due to the entirely divergent reactor concept of the RDBER: Firstly, the initial cell number per cathode area, and secondly, the hydraulic retention time in the reaction chamber (0.5 min in the flow-cells compared to 18.4 min in the RDBER). However, since in an RBC-like reactor the flow velocity over the substratum depends on the angular velocity of the substrate rather than the hydraulic residence time (HRT), we assumed that HRT was not likely to be the key process parameter that hindered biofilm growth during the preliminary cultivation experiments. In particular, because care was taken to ensure that the angular velocity of the circular blank at the outermost radius should not exceed the optimum flow velocity for cultivation of *K. spormannii* in the flow cell system. Using the circumference of the circular blanks, it was calculated that a velocity of 11 mm s^-1^ is achieved at the outermost radius at a rotational speed of 1 U min^-1^, which is within the range of the flow velocity over the cathode in the flow cell system based on the previously conducted CFD simulation (Hackbarth et al., 2020). Hence, we hypothesized, that the 44 times lower initial cell number per cathode surface ratio (3.4 * 10^9^ cm^-2^ in the flow-cells vs. 7.6 * 10^7^ cm^-2^ for the preliminary RDBER experiments) might be the limiting factor for consistent biofilm growth. Previously it was shown that at the initially uncovered cathode a part of the dissolved oxygen might be abiotically reduced into reactive oxygen species (ROS) (Hackbarth et al., 2020; Jung et al., 2020). As the initial cell number decreases, the cumulative ROS-detoxification capacity of the cells also decreases, possibly preventing a sufficient initial cathode coverage that serves as an insulator to minimize abiotic ROS formation. Therefore, we adjusted the cell number to cathode surface ratio for the subsequent experiments. Besides increasing the initial OD_600_ from 0.1 to 0.3, we also reduced the working electrode area to 700 cm^2^, in order to avoid the need of a tremendous amount of preculture, which would not have been cultivatable in our laboratory infrastructure yet. Furthermore, the current density of the system was limited by galvanometry to 1 μA cm^-2^ for 24 h (and then to 3 μA cm^-2^ for the rest of the experiment) to minimize initial ROS-formation. Operation of the RDBER under adapted process conditions now resulted in the formation of nearly continuous *K. spormannii* biofilms covering more than 90% of the cathode. Figure 3 shows a timeseries of height maps derived from OCT datasets covering a radial excerpt of the working electrode.

**Table 1:** Listing of important process parameters comparing the RDBER cultivations of *K. spormannii* performed in this study and flow cell cultivations performed in another study. LSV – Linear sweep voltammetry, CA – Chronoamperometry.

| Process parameter | Flow-cell cultivation<br>(Hackbarth et al., 2020) | Preliminary cultivation<br>(this study) | Optimized cultivation<br>(this study) |
| --- | --- | --- | --- |
| Medium | ES-medium | Es-medium | Es-medium |
| pH | 3.5 | 3.5 | 3.5 |
| Gas mixture | CO<sub>2</sub>/O<sub>2</sub> (99.5:0.5) | CO<sub>2</sub>/O<sub>2</sub> (99.5:0.5) | CO<sub>2</sub>/O<sub>2</sub> (99.5:0.5) |
| Pressure | 1.5 bar | 0.5 bar | 0.5 bar |
| Temperature | 60 °C | 60 °C | 60 °C |
| Circulation pump speed | 100 mL min<sup>-1</sup> | 500 mL min<sup>-1</sup> | 500 mL min<sup>-1</sup> |
| Volume<sub>reaction chamber</sub> | 50 mL | 8500 mL | 9850 mL |
| Hydraulic retention time | 0.5 min | 17 min | 19.7 min |
| Flow velocity / angular velocity<br>over / of substratum at r = 210 mm | up to 21 mm s<sup>-1</sup> | up to 11 mm s<sup>-1</sup> | up to 11 mm s<sup>-1</sup> |
| Electroanalytics | potentiostatic<br>LSV (first 24 h), then<br>CA (-500 mV vs.<br>SHE) | potentiostatic<br>LSV (first 24 h), then<br>CA (-500 mV vs.<br>SHE) | galvanostatic<br>1 µA cm<sup>-2</sup> (first 24 h),<br>then 3 µA cm<sup>-2</sup> |
| Volume<sub>system</sub> (incl. pipes, etc.) | 850 mL | 9500 mL | 10850 mL |
| Initial OD<sub>600</sub> | 0.1 | 0.1 | 0.3 |
| Working Electrode Surface | 20 cm<sup>2</sup> | 10000 cm<sup>2</sup> | 700 cm<sup>2</sup> |
| Initial cell number per working<br>electrode ratio | 1.34 * 10<sup>9</sup> cm<sup>-2</sup> | 3.0 * 10<sup>7</sup> cm<sup>-2</sup> | 1.47 * 10<sup>9</sup> cm<sup>-2</sup> |

**Fig. 3:**
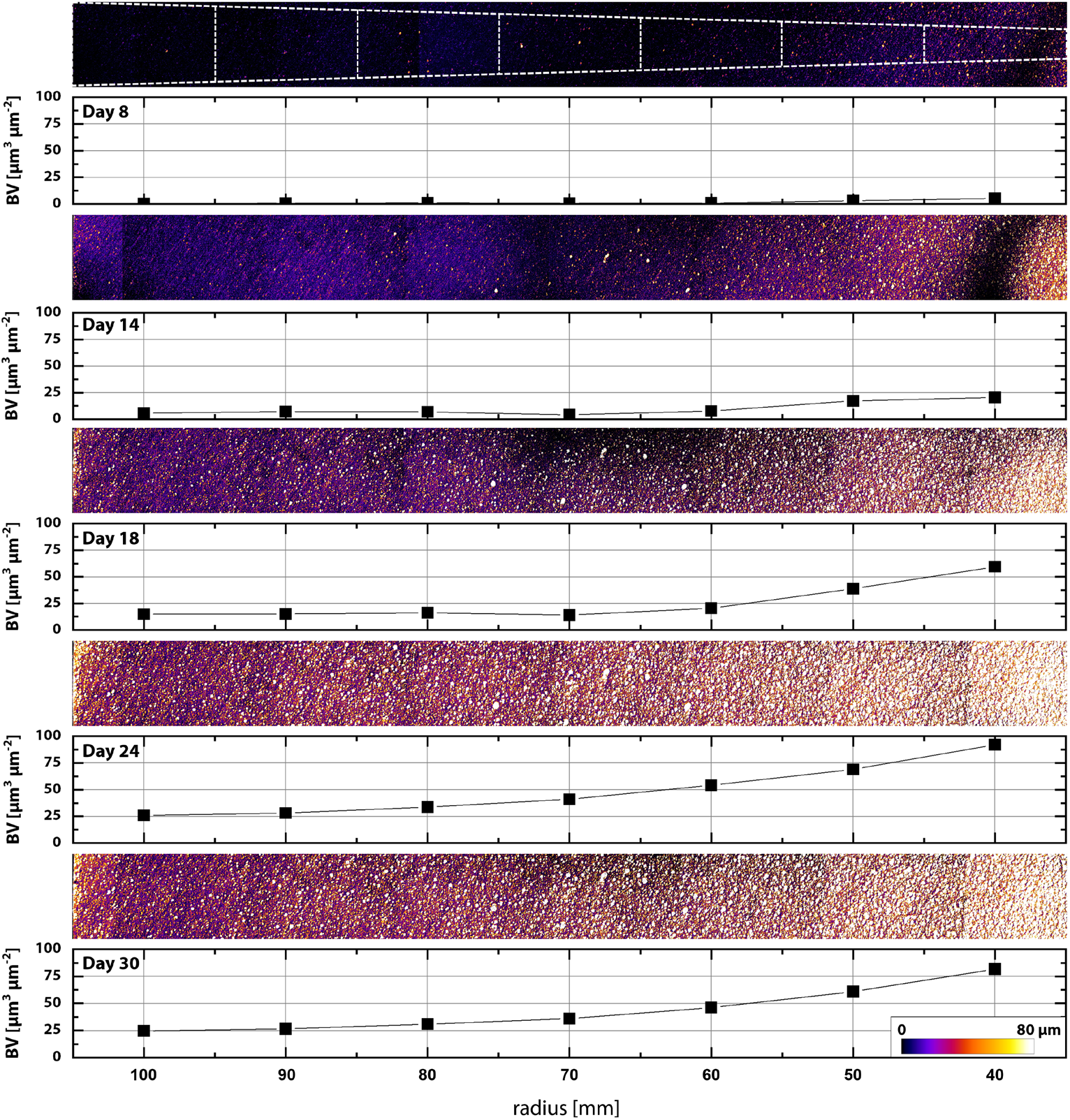
Time series of selected OCT-derived height maps of the optimized cultivation of *K. spormannii* in the 10 L-RDBER. The same radial section has been visualized on the working electrode in each case in order to show the spatial distribution of the biofilm along the rotating working electrode. Colors are coded from black (0 mm) to white (80 μm). The calculated biovolumes (*BV*) of the subtrapezoids are shown below the height profiles.

The first visible biofilm structures formed after a lag phase of 8 days near the shaft, i.e. at regions with comparatively low shear stress. A spatially resolved calculation of the local biovolume clearly shows that *BV* decreases with larger radius towards the edge of the circular blank. So, with an increased angular velocity of the substratum and therefore a higher flow velocity over the biofilm, the biovolume of the *K. spormannii* biofilm is decreasing. After 24 days *BV* reached the maximum values measured in the course of the experiment in all segments. The maximum *BV* in the outer region of the cathode averaged 25 μm^3^ μm^-2^, only about a quarter of the max *BV* of approximately 92.13 μm^3^ μm^-2^ observed in the inner visualized region. It should also be mentioned that in an abiotic control experiment, no growth-like precipitation was detected on the cathodes after 24 days. The extent to which the biovolume can be used to draw conclusions about the productivity (i.e., CO_2_-fixation rate) of the *K. spormannii* biofilm is not conclusive from this data. Nevertheless, it is known that compressed and denser biofilms generally exhibit higher turnover rates, since more catalytically active biomass is present per biovolume. At the same time, denser biofilm structures might hamper the diffusion of substrates and products within the biofilm (de Beer et al., 1994; Horn and Morgenroth, 2006), whereas the heterogeneity of the biofilm structure will also influence diffusion reaction (van den Berg et al., 2020). Certainly, the increased shear force at increased angular velocity affects the biofilm height to some extent as well. This effect may also be exploited in a future biotechnological process. Thus, a fluctuating and short-term increase of the rotational speed could serve for an optimal harvesting of the biomass from the substratum. In this way, the planktonic biomass could be partially and discontinuously extracted from the reactor, creating a self-sustaining system, since some of the detached cells can recolonize uncovered areas of the substratum. In order to compare the results of the RDBER system with those of the previously published flow cell experiments, calculated 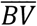 values were fitted by a logistic growth function (R^2^ = 0.98):

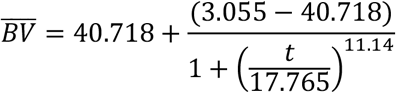

The course of the plotted values of 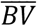 shows a significantly longer lag phase compared to the flow cell experiments. The highest biofilm accumulation rate of 6 μm^3^ μm^-2^ d^-1^ was reached between days 17 and 18 and was 45% lower compared to the flow cell experiments (see figure 6 in (Hackbarth et al., 2020)). Moreover, the maximal achieved mean biovolume in the RDBER system was 42 μm^3^ μm^-2^, which accounted to around 25% of the max. mean biovolume in the flow-cells. The highest coverage of 87% was reached on day 24, with 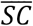 not describing a steeply increasing degressive course like in the flow-cell experiments, but a linear course until the biofilm finally detached. The biofilm in the RDBER system thus showed a generally weaker and slower growth per area of cathode surface. Notwithstanding, to our knowledge, this data shows the first successful cultivation of an aerobic, electroautotrophic organism in a BES at this scale (700 cm^-2^ working electrode). One of the reasons for the comparatively slow growth of *K. spormannii* in the RDBER was probably the moderate, limited current density (3 μA cm^-2^) - especially in contrast to the measured currents of the flow cell experiments (between 13 and 20 μA cm^-2^ at a constant working electrode potential of −500 mV vs. SHE). Although excessive ROS production does not occur if the current density is too low, the biocatalyst may not have enough electrons available for rapid and, above all, complete coverage of the substrate. Thus, in future experiments, the maximum current density should be limited to that extent only during the initial start-up phase and should also be increased as the surface coverage increases.

In addition to the limitation of the electron donor, reduced substrate availability due to lower solubility of the gaseous substrates at reduced pressure in the RDBER (0.5 bar) compared to the flow cells (1.5 bar) has to be assumed. Indeed, in their computational model of a microbial electrosynthesis process, Cabau-Peinado et al. demonstrated that CO_2_ limitations can occur and thus may play a role in low turnover rates of a MES system (Cabau-Peinado et al., 2021). According to the manufacturer, the shaft seal of the titanium shaft in the rear flange is currently the only tender spot that limits the maximum pressure in the system to 0.5 bar. Both the reactor periphery and the entire reactor jacket can be pressurized to at least 1.5 bar. Therefore, future reactor generations should incorporate a shaft seal in the rear bearing that is designed for higher pressure ranges in order to assess the impact of the pressure on the biofilm development in the RDBER. Another approach to reduce a possible substrate limitation may also consist in operating the RDBER in a similar way to an RBC. Here, the reactor would be only half filled with medium, while the gas phase - now present in the reactor chamber - would be flushed with the substrate gases. The fixed film would then move periodically through a highly concentrated CO_2_ atmosphere. Furthermore, the actual influence of the HRT in relation to the influence of the rotational speed of the working electrodes on the biofilm growth has to be investigated experimentally in order to exclude a possible growth limitation due to a too low substrate (carbon dioxide) and/or nutrient loading.

Based on these experiments, we were able to show that an RBC-like reactor is, in principle, a feasible way to upscale the previously flow cell-based cultivation of *K. spormannii* for bioplastic production from electricity and CO_2_. Since the pumping power required for successful cultivation of *K. spormannii* in an upscaled flow cell process would presumably render the overall process quickly uneconomical, an RBC-like reactor might be an interesting option to generate HRT independent crossflow velocities across the working electrode. Furthermore, the cultivation experiments of *K. spormannii* in the RDBER highlighted the crucial role of the start-up phase in aerobic MES processes. The impact of reactive oxygen species should not be underestimated in the development of future MES processes. Therefore, for the necessary development of targeted strategies, it is important to better understand the formation of reactive oxygen species at cathodes under physiological conditions, as well as the different detoxification mechanisms of electroautotrophic microorganisms.

### 3.2 Anodic operation as Microbial Electrolysis Cell

Following the successful cultivation of an electroautotrophic species in the RDBER, we used the reactor to cultivate a defined coculture of *Shewanella oneidensis* and *Geobacter sulfurreducens*. The experiments were performed as batch experiments with a full set of the 14 working electrodes (here anodes) with a total surface area of 1 m^2^. Acetate and lactate served as carbon source with an initial concentration of 20 mM each. The current density at a constant working electrode potential of 0 mV vs. SHE was recorded over the experimental course of roughly 9 days (see Fig 5). After an initial polarization phase of a few hours, characterized by negative current densities most likely due to the depletion of residual oxygen in the system, the current steadily increased until it peaked at 130 μA cm^-2^ after about 4 days. Subsequently, the current density drops again to slowly decrease to about 90 μA cm^-2^ after 9 days. Almost uniform in shape runs the graph of the gas evolution rate recorded by means of a MilliGascounter from the gas outlet of the RDBER. The hydrogen fraction in the outflowing gas was examined regularly using a MicroGC and remained almost constant over the entire experimental period (88.1 ± 1.2%). Daily liquid sampling of the reactor’s broth was conducted to determine the concentration of the organic acids via ion chromatography.

**Fig. 4:**
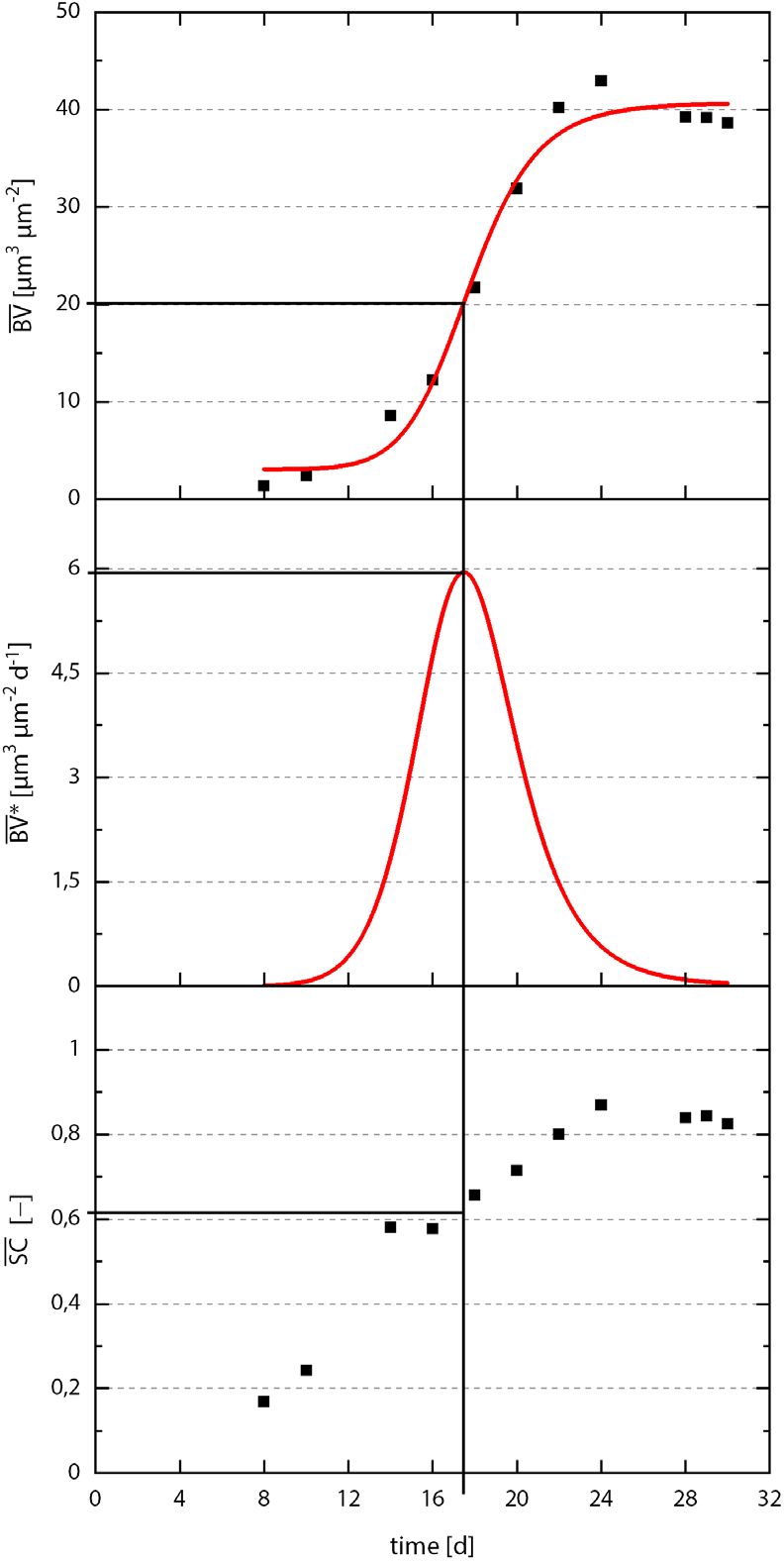
Plot of calculated biofilm parameters of the optimized cathodic operation of the RDBER over time [mean biovolume 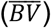, mean biofilm accumulation rate 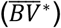, and mean substrate coverage 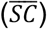].

**Fig. 5:**
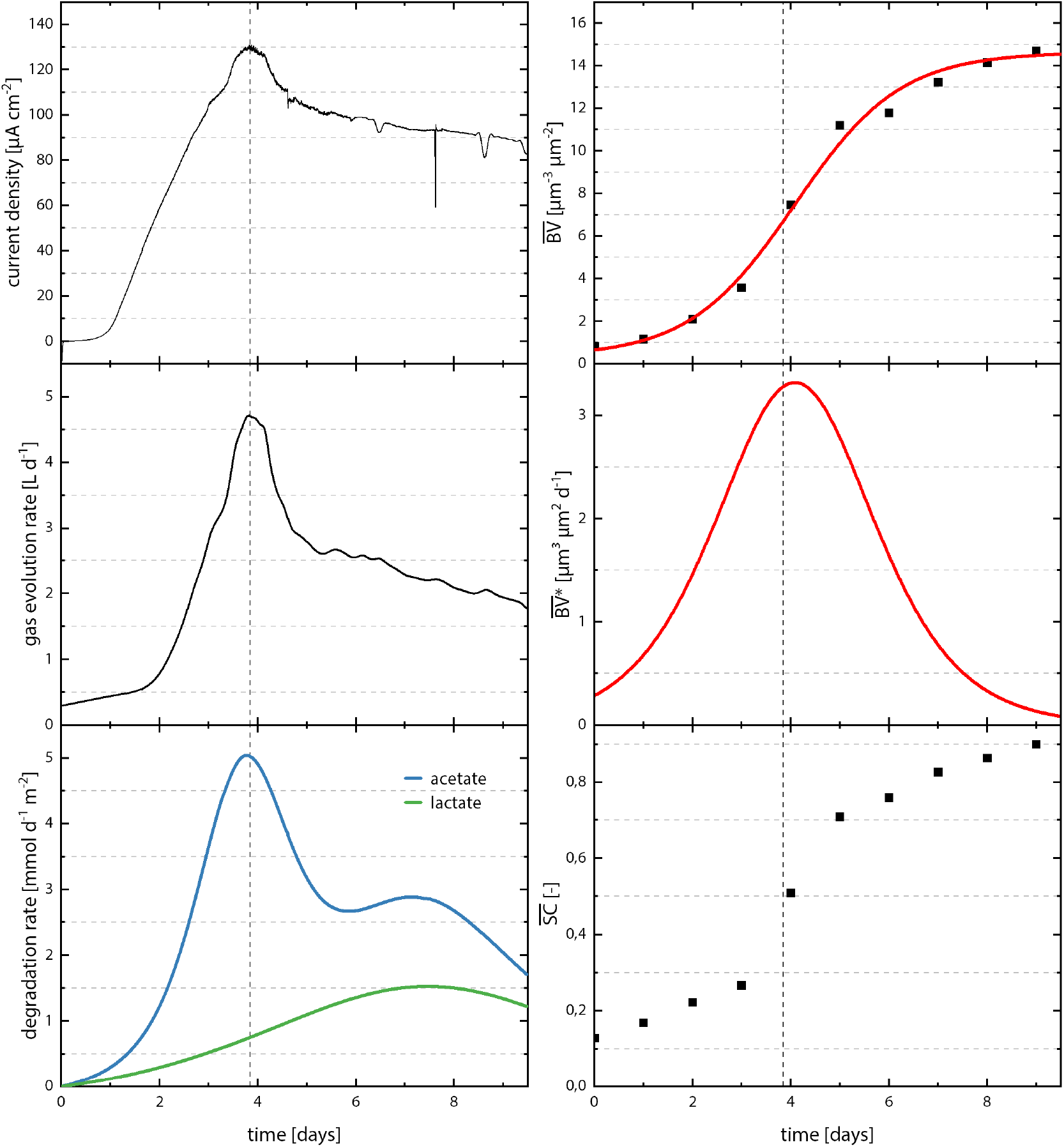
Correlation of current density, gas evolution rate and substrate specific degradation rate with calculated biofilm parameters obtained during anodic operation of the RDBER. A coculture of *Shewanella oneidensis* and *Geobacter sulfurreducens* was cultivated at a working electrode potential of 0 mV vs. SHE in a MEC-like operation. For a clearer representation, the graphs of current density and gas evolution rate were smoothed with an FFT filter. Raw data of organic acid concentrations, as well as the applied fits (displayed in color), are provided in the Supplementary Material.

The course of the concentration values over time was fitted and derived after time to obtain the specific degradation rates for acetate and lactate, respectively (please refer to Fig. S2 material for original data and fit). In this context, it was important to consider that *Shewanella* degrades one molecule of lactate into one molecule of acetate and CO_2_ or formate. Since the acetate concentration did not increase during the experiment, it was therefore assumed that the acetate produced by *Shewanella* was directly metabolized by *Geobacter*. This was taken into account when calculating the specific acetate degradation rate. Since both the peak in current data, and in the gas production rate, correlate with the peak in acetate degradation rate on day 4, these data clearly demonstrate that *Geobacter* is accounting for most of the current production, as *Shewanella* is not capable of metabolizing acetate. The maximum degradation rate for acetate was 5 mmol d^-1^ m^-2^ cathode surface and 1.5 mmol d^-1^ m^-2^ for lactate, respectively. Regular OCT scans of a radial anode section provided the basis for calculating 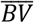 and 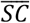. Figure 5 shows the mean biofilm accumulation rate 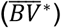 derived from a logarithmic fit through the calculated 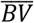, values (R^2^ = 0.99):

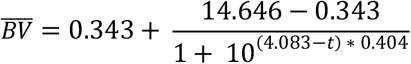

The highest biofilm accumulation rate of almost 3 μm^3^ μm^-2^ d^-1^ is likewise reached after about 4 days. Again, based on the correlations with the degradation rate, conclusions can be drawn that *Geobacter* cells appear to be the predominant species in the biofilm. This is also consistent with the literature findings that attribute rather modest biofilm capabilities to *Shewanella* compared to *Geobacter* (Engel et al., 2019). After 9 days the anodic biofilm covered around 90% of the observed working electrode surface. Interestingly, the mean surface coverage does not increase linearly as in the cathodic cultivation experiments of *K. spormannii*, but rather logarithmically. Hence, it can be concluded that the growth pattern seen in the cathodic experiments is not so much an intrinsic phenomenon of the RDBER, but probably due to the abiotic formation of ROS on uncovered cathodes described above. On the other hand, or in addition, this distinct growth pattern could also be the consequence of a synergistic effect between the two microorganisms *S. oneidensis* and *G. sulfurreducens* (Engel et al., 2019).

Comparing the performance of the RDBER with other single-chamber upscaling attempts of MECs, it is apparent on the one hand that there is still room for improvement in terms of current densities (up to 130 μA cm^-2^ in this study), but on the other hand the excellent surface to reactor volume ratio (100 m^2^ m^-3^) of the RDBER is worth emphasizing (see Table 2). Although this reached maximum current density is higher than those measured by Huang et al. or by Cotterill et al. (60 μA cm^-2^ and 79 μA cm^-2^ respectively), Wang and colleagues were able to realize tenfold higher current densities using glucose-containing medium, than those achieved in the RDBER (Cotterill et al., 2017; Huang et al., 2019; Wang et al., 2021). Furthermore, this value was surpassed again significantly by the current densities reported by Singh et al. (3400 μA cm^-2^) (Singh et al., 2021). However, when comparing current densities, it is important to keep in mind that in most MEC studies a constant voltage is applied to the anode and cathode by means of a simple power supply, whereas in our study a potentiometric control of the working electrode potential by means of a potentiostat was used to operate the RDBER. Although a potentiostat allows reproducible electrochemical conditions to be generated, the current densities achieved may be lower than with power supply operation.

**Table 2:**
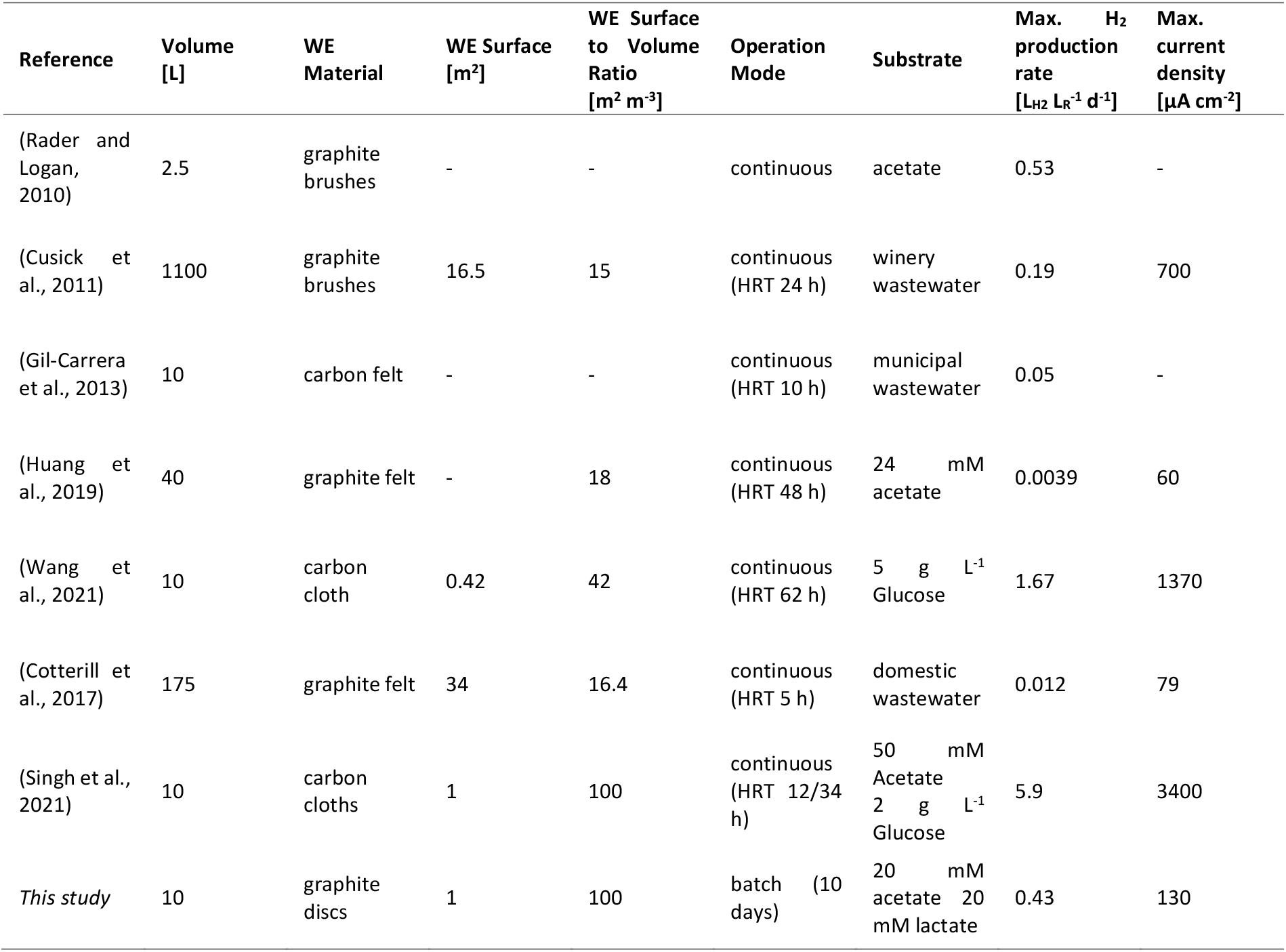
Comparison of RDBER performance with other up-scaled single-chamber MECs in the literature.

Regarding the gas production of the RDBER as another crucial criterion for assessing reactor performance, a maximum hydrogen production rate of 0.43 L L^-1^ d^-1^ (volume hydrogen per reactor volume and day) was achieved on day four. Compared to other single-chamber upscaling attempts, this value is up to several orders of magnitude higher than most of the previously achieved rates (Cotterill et al., 2017; Cusick et al., 2011; Gil-Carrera et al., 2013; Park et al., 2022). Nevertheless, it must be taken into account that some of these studies were conducted using real wastewater as substrate, which generally has lower substrate loading than the synthetic medium used in this study (Cotterill et al., 2017; Cusick et al., 2011; Gil- Carrera et al., 2013). However, under the selected process conditions, the hydrogen production rate achieved with the RDBER does not approach the rates achieved by Wang et al. and Singh et al. (1.67 and 5.9 L L^-1^ d^-1^, respectively) (Singh et al., 2021; Wang et al., 2021). Finally, when evaluating the discussed performance parameters, it should be noted that the experiments conducted here are only short-term batch tests to validate the functionality of the system. The above studies were conducted in a continuous operating mode for a significantly longer period of time, up to several months, allowing a more productive biofilm to grow on the working electrode over the longer cultivation period. In addition, the ion chromatographic data also indicate that the biofilm may have already reached a degree of substrate limitation (Fig. S2). Furthermore, it can be assumed that the optimal speed of the working electrodes will have to be determined specifically for each future application of the RDBER and thus for each specific microbial community. Although the comparability of the studies listed in Table 2 proves to be difficult to some extent, since other fundamental parameters (e.g., temperature, pH, electrode materials, electrochemical operating conditions, carbon source and loading rate, HRT, and the microbial community) may differ significantly in addition to the reactor system itself, it nevertheless appears that the RDBER - especially if reactor performance is further enhanced - can compete quite well alongside with existing MEC reactor concepts.

## 4. Conclusions

In conclusion, the study aimed on the design of a novel, scalable electrochemical reactor concept that meets the requirements of biotechnological pure culture cultivations. During the construction of the single-chamber system, emphasis was laid on a long operating lifetime and low-maintenance operation. The successful cultivation of different electroactive model organisms, both in anodic and cathodic operation, prove on the one hand the basic functionality of the system. On the other hand, we were able to demonstrate that an increase in the working electrode area is not only possible by numbering up individual small reactor systems, as is often the case for METs. Although the experiments discussed here suggest that the RDBER might be a viable reactor concept, in particular for specific niche processes like the aerobic, microbial electrosynthesis, there should be no doubt that more data from long-term studies under optimized and continuous operating conditions are needed to reliably evaluate the performance of the RDBER compared to other reactor concepts. However, we think that if microbial electrochemical technologies are to become truly competitive, a greater range of creative reactor designs that can be scaled up without excessive numbering-up is necessary. A further scale-up of the RDBER is conceivable as well: While the diameter of the electrode discs is limited mainly by the angular velocity at the outer edge, the shaft can be extended (and equipped with more electrodes) until it is unable to statically support its own weight.

## Acknowledgements

Special thanks go to Prof. Harald Horn for the initial idea behind the reactor concept, to Axel Heidt as the person responsible for the IC measurements and to Erwin Wachter, Joachim Lang and Alfred Herbst from the metal workshop of the EBI for their work and support during the construction of the reactor.

## CRediT authorship contribution statement

**Max Hackbarth:** Investigation, Conceptualization, Methodology, Visualization. **Johannes Gescher:** Conceptualization, Writing - Review & Editing, Funding acquisition. **Harald Horn:** Conceptualization, Writing - Review & Editing, Funding acquisition. **Johannes Eberhard Reiner:** Investigation, Conceptualization, Methodology, Visualization, Writing - Original Draft & Editing.

