## Supplementary Material for "A scalable, Rotating Disc Bioelectrochemical Reactor (RDBER) suitable for the cultivation of both cathodic and anodic biofilms"

**Supplementary Figure 1**

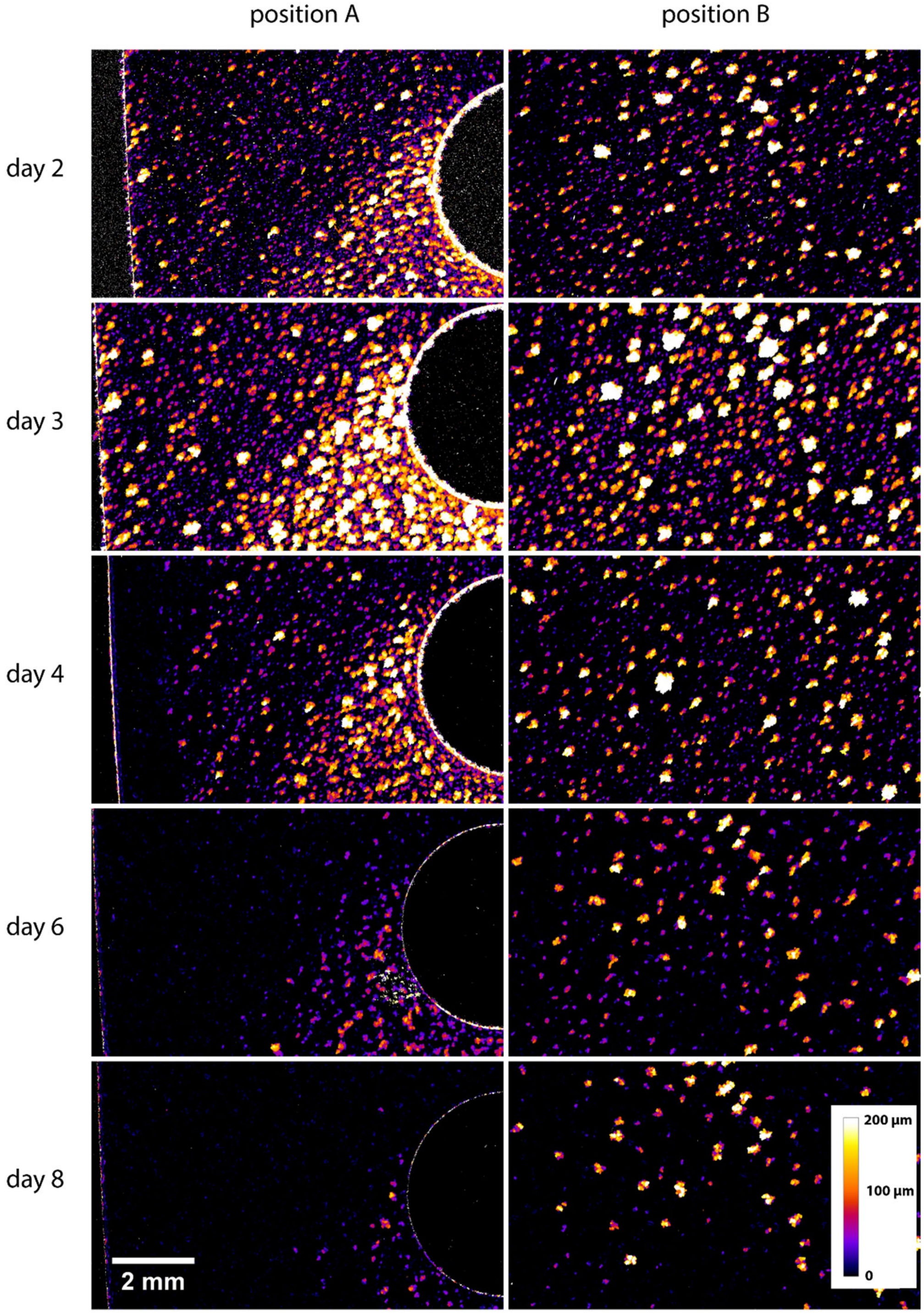


Supplementary Fig. S1: Time series of height maps calculated from OCT data sets taken from two spots on a radial stripe of the rotating cathode. Position A shows a location at the outer edge of the graphite slice, whereas position almost is at the inner visualizable edge.

**Supplementary Figure 2**

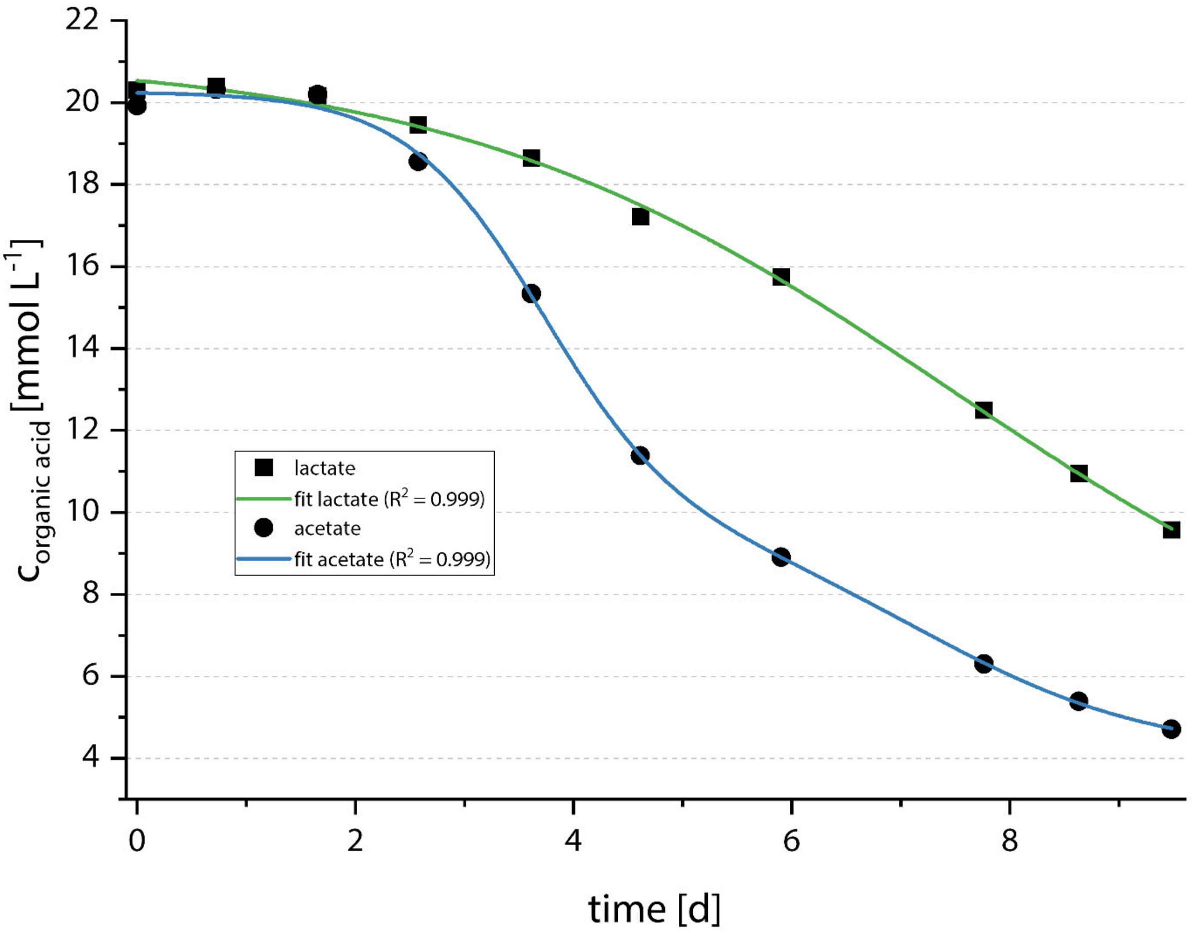


Supplementary Fig. S2: Original IC data on the organic acid degradation during anodic operation of the RDBER. The green and the red line correspond to the fitted functions used for calculation of the specific degradation rates.
